# *Chloris distichophylla* Lag. (Poaceae) as stink bug (Pentatomidae) shelter in soybean and corn off-season in southern Brazil

**DOI:** 10.1101/670943

**Authors:** Eduardo Engel, Mauricio P. B. Pasini, Daniele C. Hörz, Rafael P. Bortolotto

## Abstract

The objective of this study was to evaluate the composition and abundance of dormant bedbugs in *Chloris distichophylla* Lag (Poales: Poaceae) over the soybean and corn off-season. The work was carried out in the municipality of Cruz Alta, over the soybean and corn off-season between 2014 and 2018. Plants of *C. distichophylla* with different clump diameter were sampled, and the bugs contained in them were counted and submitted to data analysis for the evaluation of the composition, structure, and diversity of occurring species. At the end of the experiment, 3543 hibernating adults were counted and divided into six species: *Euschistus heros* (F.), *Dichelops furcatus* (F.), *Dichelops melacanthus* Dallas, *Edessa meditabunda* (F.), *Edessa ruformaginata* (De Geer) (Hemiptera: Pentatomidae). The species *E. heros* was the most abundant, followed by *D. furcatus*. The diameter of the clumps directly affects the population density of the stink bugs. Finally, *C. distichophylla* is shown as a hibernate favorable to the maintenance of the stink bug populations over the soybean and corn off-season.

## Introduction

In Brazil, soybean and corn crops are of great importance to its economy, since they, directly and indirectly, generate employment and income (Brazil 2018). Regarding the maintenance of the productivity of both crops, it is necessary to know about the behavior of insects considered to be pests of economic importance (Abrol 2013).

Among the main pests of soybean and corn, stink bugs are highlighted due to their capacity of damage (Panizzi *et al*., 2012). In the soybean crop, these insects, directly and indirectly, generate losses through sap sucking, therefore reducing grain weight, lowering the physiological quality of seeds besides causing pod abortion (McPherson 2018). In corn, its damage is significant at the beginning of development, where it causes a reduction in the population stand of the plants using the so-called “leaf rolling” of the plant, making its development impossible (Gassen 1996).

In addition to crop damage, another factor that determines the adaptive success of these insects is their ability to search for alternative hosts during the crop off-season, either to their feeding and completion of their cycle or to hibernate (Panizzi 1997, Klein *et al* 2012, Smaniotto & Panizzi 2015).

Pasini et al. (2015) observed the effect of different cropping systems on the population density of stink bugs and found that the soybean-rice succession was beneficial for the control of the *Tibraca limbativentris* Stal. (Hemiptera: Pentatomidae). On the other hand, Pasini *et al*. (2018) verified the dependence of the *T. limbativentris* bugs on host plants during the off-season in an irrigated rice field. In this context, the knowledge about the plants associated with these insects during the crop off-season becomes of extreme importance for integrated pest management. For Link & Grazia (1987), the knowledge of the host plants helps in the studies of ecology, management, and prediction of species that cause harm to the cropped plants.

In this context, the objective of this study was to determine the diversity and abundance of stink bug species associated with *Chloris distichophylla* Lag (Poales: Poaceae) plants near the soybean-corn succession area during the off-season.

## Material and methods

The experiment was conducted at the University of Cruz Alta, Cruz Alta, Rio Grande do Sul state, (Time Zone 22, 244138; 6835737 UTM). According to the classification of Köppen, the climate in the region is classified as Cfa, with average temperature less than 18°C (mesothermic) in the coldest month, and average temperature higher than 22°C in the hottest month, with hot summers, occasional and tendency of rainfall concentration in summer; however with no defined dry season (Kuinchtner & Buriol 2016) during the off-season in the cropped area in soybean-corn succession in 2014, 2015, 2016, 2017 and 2018.

Plants with 5, 10, 15, 20, and 25 centimeters of clump diameter were sampled. The individuals occurring inside the clumps were visually counted, for the unidentified insects, which were taken to the Laboratory of Entomology at the University of Cruz Alta for identification. For each clump diameter, ten plants per year were sampled with a maximum distance of 15 meters from the edge of the cultivation area and a minimum distance of 20 meters between the evaluated host plants. Each plant was considered an independent experimental unit, totaling 250 experimental units at the end of the five years.

In order to determine the effect of clump diameter on population density, linear regression analysis was used. Quantitative data were analyzed through the alpha diversity (diversity and distribution of species abundance - SAD). The suitability of SAD was tested in four models: geometric, broken-stick, log-series, and log-normal. The sampling sufficiency curve for the abundance of canola-associated pentatomids was obtained with 999 randomizations and compared with the non-parametric richness estimators Chao 1, Chao 2, Jackknife 1 and Jacknife 2 to determine the sampling efficiency, according to the methodology used by Bianchi *et al*. 2018. Each richness estimator takes into account a different parameter, that is, the occurrence of singletons, doubletons, unique, and duplicates. All analyses were performed using the software PASt 3.34 (Hammer *et al*. 2001).

## Results

A total of 3543 individuals belonging to the six species were counted (Table 1). Among the species of the sampled stink bugs, the greatest variability in the number of insects was found for species *Euschistus heros* F. followed by *Dichelops furcatus* F. (Fig 1).

**Table 1.** Composition, abundance (N) and frequency (F) of stink bug (Pentatomidae) sampled in *Chloris distichophylla* Lag. (Poales: Poacae) over soybean and corn off-season from 2014 to 2018 in Cruz Alta, Rio Grande do Sul, Brazil.

| Species | N | F (%) |
| --- | --- | --- |
| *Euschistus heros* (F.) | 1837 | 52 |
| *Dichelops furcatus* (F.) | 1019 | 29 |
| *Dichelops melacanthus* (Dallas) | 33 | 1 |
| *Edessa meditabunda* (F.) | 467 | 13 |
| *Edessa ruformarginata* (De Geer) | 27 | 1 |
| *Piezodorus guildinii* (Westwood) | 160 | 5 |
| Total | 3543 | 100 |

**Fig 1.**
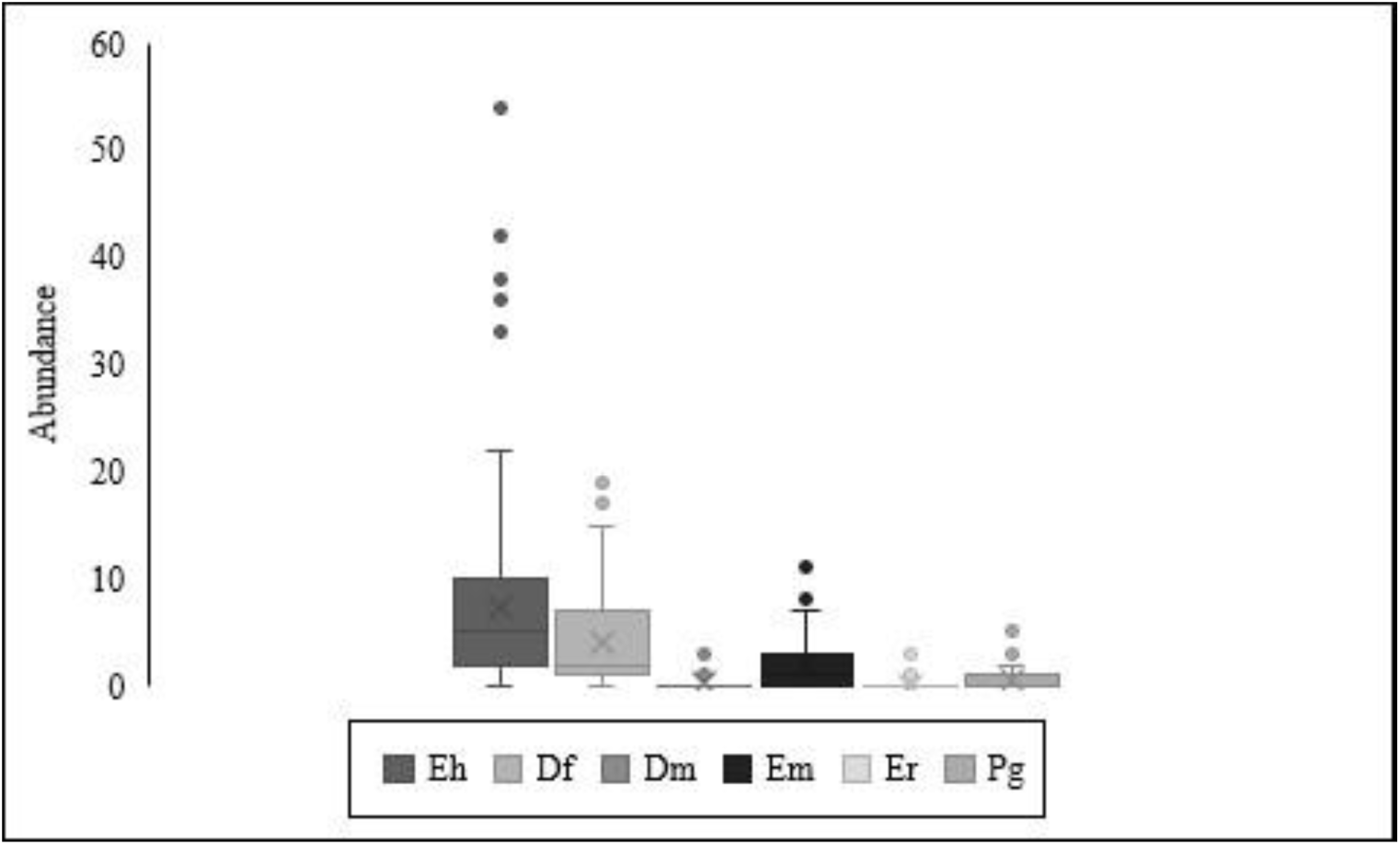
Box plot of the number of stink bugs sampled in *Chloris distichophylla* Lag (Poales: Poacaeae) during the soybean and corn off-season from 2014 to 2018. Cruz Alta, Rio Grande do Sul. Eh (*Euschistus heros*), Df (*Dichelops furcatus*), Dm (*Dichelops melacanthus*), Em (*Edessa meditabunda*), Er (*Edessa ruformaginata*) and Pg (*Piezodorus guildinii*).

It was found in this study that at least four out of the six species captured are harmful to soybean and corn crops (except only *E. meditabunda* and *E. ruformaginata*). The most abundant species were *E. heros* (N = 1837), *D. furcatus* (N = 1019) and *E. meditabunda* (N = 467) (Table 1). The sum of these species corresponded to 94% of the individuals sampled (Table 1). For the analysis of the abundance distributions among species, it was found a significance for the geometric model (k = 0.6044; chi^2 = 147.3; p < 0.0001) (Fig. 2).

**Fig 2.**
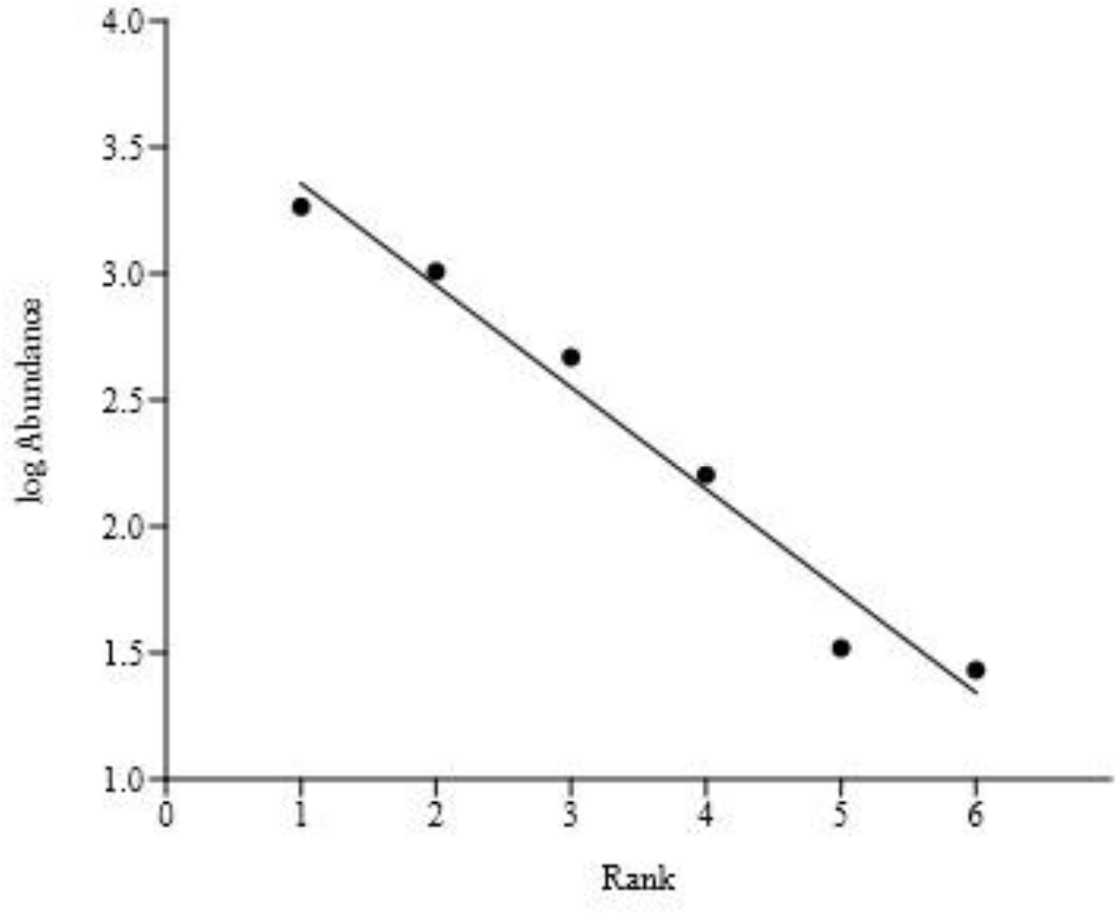
Distribution of the abundances of pentatomid bugs sampled in *Chloris distichophylla* Lag. (Poales: Poaceae) in the soybean and corn off-season crop between 2014 and 2018. Cruz Alta, Rio Grande do Sul, Brazil.

A trend towards asymptote from the thirtieth sample was observed through the rarefaction curve. This trend suggests a stabilization of the number of species to be sampled, indicating sample sufficiency. For all species richness estimators (Chao 1, Chao 2, Jacknife 1 and Jacknife 2), a value equal to that observed, reinforcing the idea of sufficiency in species sampling in the evaluated plants (Fig 3).

**Fig 3.**
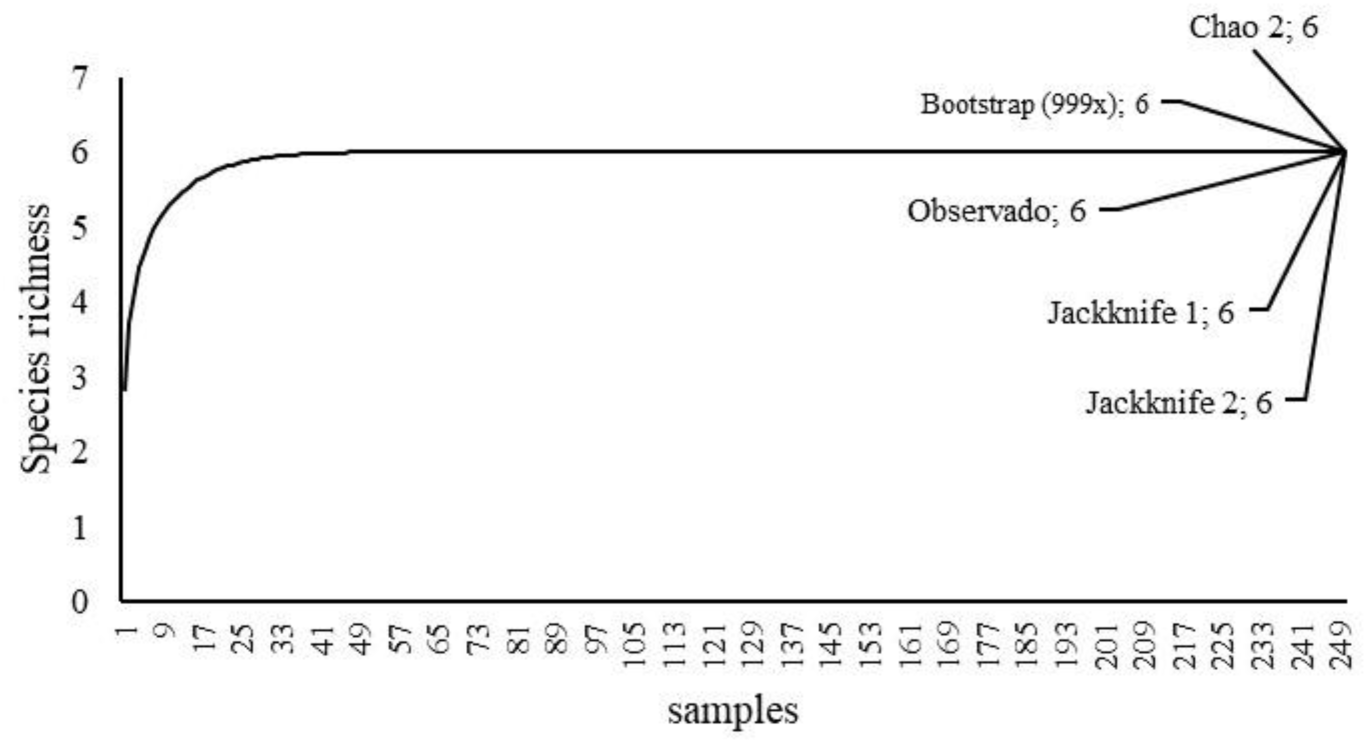
Rarefaction curve and species richness estimators for pentatomid bugs sampled in *Chloris distichophylla* Lag. (Poales: Poaeceae) during the soybean and corn off-season from 2014 to 2018. Cruz Alta, Rio Grande do Sul, Brazil.

A direct effect of the *C. distichophylla* clump diameter was observed on the population density of the sampled bugs (Fig. 4). This direct effect may be associated with competition for space (Klein et al., 2012), but there is also temperature regulation within the clumps (Dennis *et al*. 1994).

**Fig 4.**
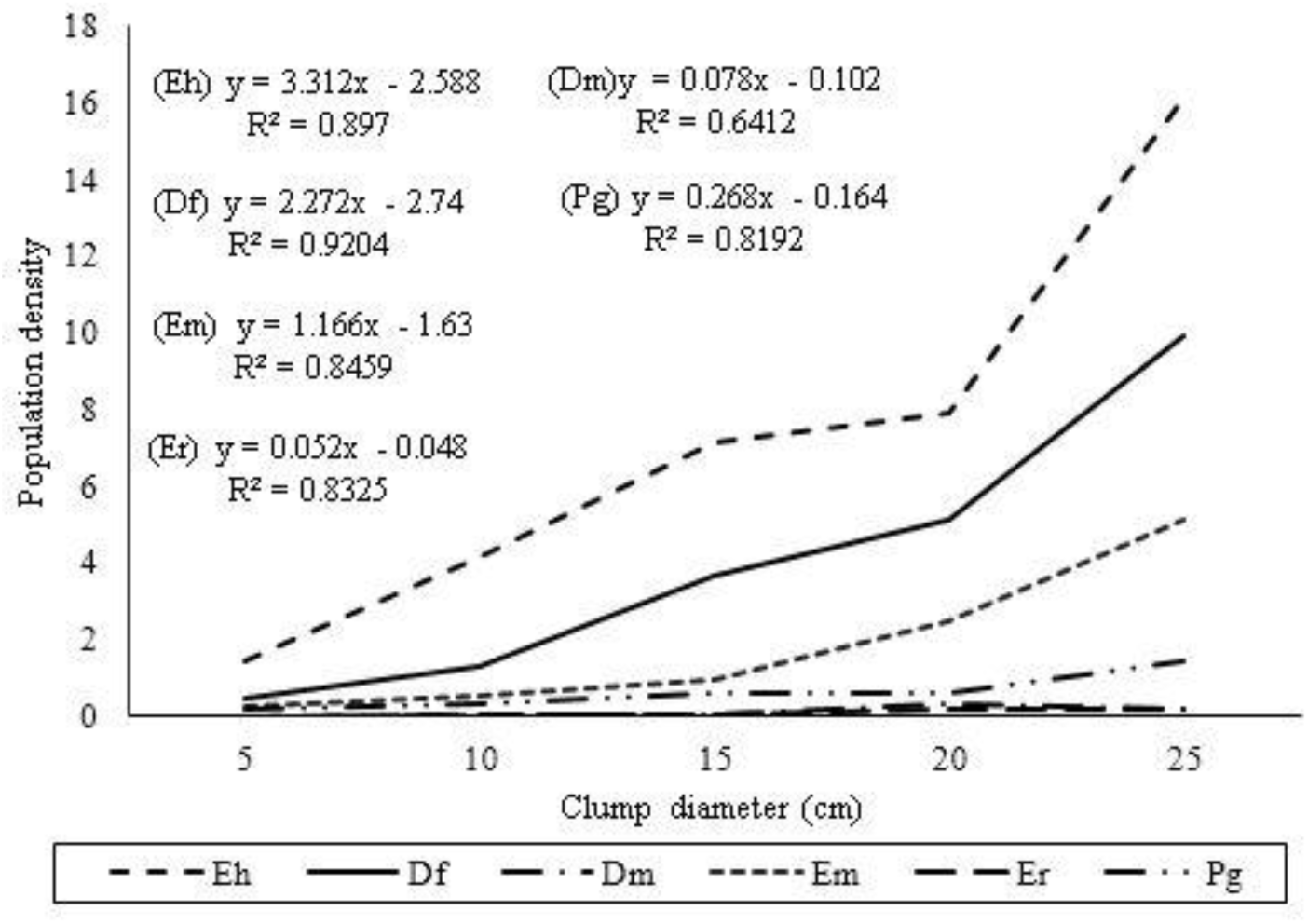
Population density of pentatomid bugs as a function of the clump diameter of *Chloris distichophylla* Lag. (Poales: Poaceae) during the soybean and corn off-season. Cruz Alta, Rio Grande do Sul, Brazil. Eh (*Euschistus heros*), Df (*Dichelops furcatus*), Dm (*Dichelops melacanthus*), Em (*Edessa meditabunda*), Er (*Edessa ruformaginata*) and Pg (*Piezodorus guildinii*).

No nymphs were observed among the sampled individuals over the whole monitoring period, therefore, indicating no feeding and oviposition preference for the plant in question. Thus, its occupation was only for survival during the crop off-season. The same situation was observed in *A. bicornis* (Klein *et al*. 2012, Pasini *et al*. 2018, Engel *et al*. 2018).

## Discussion

Because of the large number of host and associated plants, the economic importance of stink bugs is related not only to their feeding behavior but also to their forms of survival (Smaniotto & Panizzi 2015, Pasini *et al*. 2018). Their wide geographical distribution makes them pests not only in Brazil, but in the world, either in the form of invasive or endemic species (McPherson 2018).

The knowledge of the different forms of survival and plants associated with pentatomid bugs during the off-season allows the optimization of the management by identifying areas of control and infestation (Engel et al. 2018). This work identified the diversity of a stink bug assembly associated with *Chloris distichophylla* located in the border of the cultivation area during the off-season of soybean and corn crops for five consecutive years.

Several studies highlight the ways of survival of stink bugs in unfavorable periods. Medeiros & Megier (2009) found associated plants for *E. heros*, which served as a food source during the off-season. Klein *et al* (2012) identified *Andropogon bicornis* L. (Poaceae) as a hibernacle of a series of stink bugs in off-season of rice crop. The same situation was observed by Pasini *et al*. (2018), who also observed the occurrence of *T. limbativentris* in *Andropogon lateralis* (Poaceae).

A series of species considered of economic importance was observed for *C. distichophylla*; however, extensive dominance was observed for *E. heros, D. furcatus*, a factor explained by the crop system in succession with the soybean and corn crops, favoring the development and permanence of these insects in the cropping area.

It is also verified the influence that the clump diameter of the associated plants during the off-season has on the population density. Concerning *C. distichophylla*, a higher population density due to clumps with larger diameter was found. Pasini *et al*. (2015) emphasize as factors related to the selection of insects for plants, the structural complexity and their distance of the border to the area of cultivation, a fact evidenced for the *T. limbativentris* (Klein *et al*. 2012, Pasini *et al*. 2015, Pasini *et al*. 2018).

The volume of plant mass in associated plants may be related to the capacity of maintenance of the microclimatic conditions in clumps closest to the minimum temperature required by pentatomid bugs. Therefore, plants with larger diameters of clump allow a higher survival chance for these insects (Dennis *et al*. 1994, Howe & Jander 2008).

Considering the species of stink bugs found in the study, mainly those of economic importance (*E. heros, D. furcatus, D. melacanthus*, and *P. guildinii*), it was verified the importance of *C. distichophylla* plants for the maintenance of populations during the soybean and corn off-season. As a result, methods of management and sampling of this environment can be adopted in order to contribute to the population suppression of these organisms.

